# Disrupted reinforcement learning during post-error slowing in ADHD

**DOI:** 10.1101/449975

**Authors:** Andre Chevrier, Mehereen Bhaijiwala, Jonathan Lipszyc, Douglas Cheyne, Simon Graham, Russell Schachar

## Abstract

ADHD is associated with altered dopamine regulated reinforcement learning on prediction errors. Despite evidence of categorically altered error processing in ADHD, neuroimaging advances have largely investigated models of normal reinforcement learning in greater detail. Further, although reinforcement leaning critically relies on ventral striatum exerting error magnitude related thresholding influences on substantia nigra (SN) and dorsal striatum, these thresholding influences have never been identified with neuroimaging. To identify such thresholding influences, we propose that error magnitude related activities must first be separated from opposite activities in overlapping neural regions during error detection. Here we separate error detection from magnitude related adjustment (post-error slowing) during inhibition errors in the stop signal task in typically developing (TD) and ADHD adolescents using fMRI. In TD, we predicted that: 1) deactivation of dorsal striatum on error detection interrupts ongoing processing, and should be proportional to right frontoparietal response phase activity that has been observed in the SST; 2) deactivation of ventral striatum on post-error slowing exerts thresholding influences on, and should be proportional to activity in dorsal striatum. In ADHD, we predicted that ventral striatum would instead correlate with heightened amygdala responses to errors. We found deactivation of dorsal striatum on error detection correlated with response-phase activity in both groups. In TD, post-error slowing deactivation of ventral striatum correlated with activation of dorsal striatum. In ADHD, ventral striatum correlated with heightened amygdala activity. Further, heightened activities in locus coeruleus (norepinephrine), raphe nucleus (serotonin) and medial septal nuclei (acetylcholine), which all compete for control of DA, and are altered in ADHD, exhibited altered correlations with SN. All correlations in TD were replicated in healthy adults. Results in TD are consistent with dopamine regulated reinforcement learning on post-error slowing. In ADHD, results are consistent with heightened activities in the amygdala and non-dopaminergic neurotransmitter nuclei preventing reinforcement learning.

## 1 Introduction

Attention-deficit/hyperactivity disorder (ADHD) is a common neurodevelopmental disorder of childhood associated with distinctive reinforcement learning evident in altered behavior and neural responses on prediction errors (Durston et al., 2007; Furukawa et al., 2014; Ma et al., 2016; Plichta and Scheres, 2014). Reinforcement learning adjusts behavior in proportion to prediction error magnitude, defined as the difference between the actual and expected value of an outcome. Prediction errors, regulated by midbrain dopamine (DA) neurons (Ljungberg et al., 1991; Schultz, 2016a; Waelti et al., 2001), consist of an initial error detection stage independent of error magnitude, followed by processing related to error magnitude (Schultz, 2016b). In the ventral striatum, activity related to error magnitude adjusts task-related networks by influencing the threshold for the passage of task-related activity through the dorsal striatum (Horvitz, 2002). Thresholding influences from ventral to dorsal striatum are carried by both direct projections and indirect projections via substantia nigra (SN). Given that reinforcement learning is critically dependent on DA regulated striatal thresholding, it is essential to determine the gating conditions in the striatum during altered error processing in ADHD. However, DA regulated thresholding influences from ventral to dorsal striatum have not been studied with neuroimaging, or in any context other than invasive experiments in animals.

We propose that thresholding influences from ventral to dorsal striatum have not previously been identified with neuroimaging because they have been combined with opposite activities in the same regions during error detection. In a previous study of errors in the stop signal task (SST), we separated error detection from error magnitude related adjustment activity for the first time using functional magnetic resonance imaging (fMRI) (Chevrier and Schachar, 2010). The SST is a widely used measure of response inhibition and error detection and allows for separation of error detection from error magnitude related adjustment. The SST presents choice response stimuli that occasionally (33%) must be stopped (Logan et al., 1984). The stop signal delay adapts to performance so that only half of these trials can be stopped on average. Unsuccessful stop trials are negative instrumental prediction errors, which should be followed by adjustments proportional to error magnitude just as in reward tasks (Gallistel and Gibbon, 2000).

The nature of the SST is such that slowing is the only form of adjustment available to errors of greater magnitude, affording a minimally complex imaging design capable of separating response, error detection and adjustment-related stages of activity. Using this approach in a previous study of healthy adults, we found that dorsal striatum deactivated on error detection, which we proposed interrupts ongoing processing by withdrawing subcortical support from right frontoparietal regions that activate during response phases (Chevrier et al., 2015, 2007; Chikazoe et al., 2009) when errors are detected. We also found deactivation of ventral striatum on errors followed by greater than median post-error response times, which we proposed exerts error magnitude related thresholding influences on SN and dorsal striatum necessary for reinforcement learning. The need to separate error detection from magnitude related adjustment in order to identify their functional roles in interrupting ongoing processing and striatal thresholding is apparent from the fact that post-error slowing activated the same regions of dorsal striatum and SN that deactivated on error detection.

Here we use our previous imaging approach to compare typically developing (TD) adolescents and adolescents with ADHD. In TD, we attempt to replicate our previous findings of deactivations in dorsal and ventral striatum on error detection and post-error slowing, and perform intersubject correlation analyses on these activities to confirm our hypotheses about their functional roles in interrupting and adjusting task related processing

Firstly, if deactivation of dorsal striatum interrupts task-related processing when subjects detect that they have made an erroneous response, then subjects who activate task-related networks more strongly during response-phases should require greater deactivation of dorsal striatum when errors are detected. Therefore, we predict that deactivation of dorsal striatum on error detection should be correlated with right frontoparietal response phase activities.

Secondly, if deactivation of ventral striatum on post-error slowing exerts error magnitude related thresholding influences on dorsal striatum, then subjects with greater deactivation of ventral striatum on post-error slowing should also have greater activity in SN and dorsal striatum; Therefore, we predict that deactivation of ventral striatum should be correlated with activity in SN and dorsal striatum during post-error slowing.

In ADHD, striatal thresholding could be disrupted by heightened activity in the amygdala, which encodes prediction errors in response to aversive stimuli (McHugh et al., 2014). Amygdala responses to aversive stimuli are exacerbated in ADHD and are correlated with affective symptoms (Maier et al., 2014). Here we compare TD and ADHD to determine whether heightened amygdala responses to prediction errors observed in previous studies of ADHD that have used more emotional stimuli such as facial expressions, also occur on instrumental errors in the SST. Heightened amygdala activity could disrupt reinforcement learning because the level of amygdala input to the ventral striatum has a specific role in regulating striatal thresholding influences by affecting lateral connections from ventral to dorsal striatum known as the striatonigrostriatal system (SNS) (Shiflett and Balleine, 2010).

The lateral connections of the SNS cause the level of activity in more ventral parts of the striatum to affect the threshold for activity in more dorsal parts of the striatum (Haber et al., 2000). Elevated activity in the ventral striatum, such as from heightened amygdala input, therefore causes increased thresholds on the passage of cognitive activities through more dorsal parts of the striatum, referred to as limbic motor interfacing (Mogenson et al., 1980; Morrison et al., 2017). Limbic motor interfacing would thus render the striatum incapable of sustaining error magnitude related thresholding influences in dorsal striatum. Therefore, we predict that altered thresholding function caused by heightened amygdala activity would be evident in correlation of ventral striatum with the amygdala and not with dorsal striatum and SN.

DA activity that supports thresholding function can also be influenced by other neurotransmitter systems like norepinephrine (NE), serotonin (5HT) and acetylcholine (Ach). All of these neurotransmitter systems respond to prediction errors (Clewett et al., 2014; Cohen et al., 2015; Diaconescu et al., 2017; Payzan-LeNestour et al., 2013), interact with DA via direct and indirect pathways (Boureau and Dayan, 2011; Yu and Dayan, 2003), and exhibit altered functioning in ADHD (Bidwell et al., 2011). We inspected error detection and post-error slowing maps for heightened activity in raphe nucleus (5HT), locus coeruleus (LC) (NE), and basal forebrain/medial septal nuclei (Ach) in ADHD compared to TD. Heightened responses in these nuclei could influence DA regulation of reinforcement learning by affecting activity in SN as well as influencing striatal thresholding by affecting activity in the amygdala. However, the effects of non dopaminergic neurotransmitter nuclei on DA regulated thresholding have not previously been studied. Here we test for the presence of such interactions using significant post-error slowing activities and group differences in neurotransmitter nuclei as seeds for intersubject correlation analyses. These correlation maps were inspected for the presence of whole brain corrected correlations with SN, the amygdala and with one another. If post-error slowing generates an altered competition for control of DA in ADHD, then non dopaminergic neurotransmitter nuclei should exhibit heightened activities and altered correlations compared to TD.

In order to validate our results, we performed whole brain confirmatory correlation analyses in SN and raphe nucleus where whole brain patterns of connectivity have been well characterized, and compared results from TD with an independent replication sample of healthy young adults.

## 2 Materials and methods

### 2.1 Subjects

14 adolescents diagnosed with ADHD (7 male) and 14 TD adolescents (9 male, 9-18 years) were included in this study. Subjects gave informed, written consent and the study was approved by the Hospital for Sick Children institutional research ethics board. Written informed consent was obtained from the parents of all participants under the age of 16. ADHD subjects on stimulant medication (n = 6) were asked to stop administration 24 hours prior to the scan to eliminate drug-induced BOLD changes (Dodds et al., 2008).

Subjects and their parents were interviewed separately and together using the parent interview for child symptoms (PICS-IV (Ickowicz et al., 2006)). Intelligence was assessed using the Wechsler Intelligence Scale for Children (WISC-IV). ADHD subjects met diagnostic and statistical manual of mental disorders (DSM-5) criteria for ADHD (at least six out of nine inattentive symptoms, hyperactive-impulsive symptoms, or both according to at least two of three informants (parents, teacher and/or patient self-report). ADHD subjects also showed moderate to severe impairment in both school and home settings (Global Assessment Scale (Shaffer, 1983) score < 60). Subjects were excluded if they had any comorbid psychiatric or neurological disorder other than oppositional defiant disorder (ODD) or learning disability within the previous 12 months (e.g., obsessive compulsive disorder, Tourette syndrome, major depressive, anxiety or pervasive developmental disorder), an IQ score of below 80 on verbal and performance scales or any medical issues that would impact fMRI participation. Subjects with contraindications for MRI (metal braces or metal fragments in their body) were also excluded.

Nine ADHD subjects were diagnosed with ADHD combined subtype, five met criteria for inattentive subtype, and two also met DSM-5 criteria for ODD. Control subjects were assessed in a comparable manner and reported no psychiatric or medical disorders. All subjects were right-handed and had normal vision and hearing.

### 2.2 Behavioral task

The stop signal task (SST) (Logan et al., 1997) involves a primary choice reaction time task and a secondary stop task. Trials began with a fixation point in the centre of a black screen (500 ms), followed by the go-stimulus (1000 ms). Subjects were instructed to respond as quickly and accurately as possible with their left thumb when the letter “X” appeared or with their right thumb when the letter “O” appeared. In 33% of trials, a stop signal (background colour change from black to red) followed the go stimulus. Subjects were instructed to stop if they saw the stop signal, but not to wait for stop signals. The initial stop signal delay was 250 ms and increased/decreased by 50 ms after successful/unsuccessful stop trials, ensuring 50% stop errors on average.

The task involved 224 trials, requiring a total scan time of 15 minutes. Therefore, each subject contributed 37.3 errors on average, approximately half of which (18.7) are followed by greater than median post-error slowing. The appropriate contrast to noise measure for event-related detection sensitivity is dependent on the variance inherent to the BOLD response, which should rely on the number of events in a given design. However, the close range of contrast to noise measurements across designs with varying numbers of events shows that contrast to noise is not sensitive to the low number of post-error slowing events in the current (Welvaert and Rosseel, 2013).

Inter-trial interval (ITI) was jittered to maximize the number of independent equations in the deconvolution analysis, using trials of 2.5 or 3.5 seconds. Every fourteenth trial was followed by a 17.5 second rest (blank screen). Two short and one long but rare intertrial interval as used here is optimal for separating within-trial activities (Ollinger et al., 2001). Trial order was pseudo randomized so that the current trial did not predict the subsequent type of trial. Mean go response time (RT) was observable from the 67% of trials in which no stop signal appeared. Stop signal reaction time (SSRT) was estimated by subtracting the mean delay on stop signal trials from the mean RT on trials with no stop signal. Behavioral scores (within-group means and between-group differences) were analyzed using two-tailed t-tests.

### 2.3 Scanning Parameters

Imaging was done with a GE LX 1.5T MRI scanner (GE Healthcare, Waukesha, WI). Anatomical data were acquired with a standard high-quality SPGR sequence (120 slices, 1.5 mm thick, FOV 24 cm, 256 × 256 matrix). Functional data were collected using a GRE-EPI sequence with an 8-channel head coil (TE = 40; TR = 2,000; Flip angle =90 degrees; 24 slices; 6 mm thick; FOV 24 cm; 100 kHz readout bandwidth; 64 × 64 in-plane resolution). Behavioral data were collected using a fiber-optic response system interfaced to a laptop running the SST.

### 2.4 Single subject analysis

Functional data were analyzed using AFNI version 16.0.09 (Cox, 1996). Images were motion corrected and inspected to ensure motion did not exceed 3 mm or 3 degrees. We used a standard motion correction algorithm and censored noisy time points (>3.5 median absolute deviations). We used a general linear model of stimulus vectors convolved with the hemodynamic response function (HRF) using AFNI’s 3dDeconvolve program. Estimates of baseline (seventh order polynomial) were generated along with 6-point HRF’s for all event types (HRF delay of 2 time points (TR = 2s), followed by HRF window lasting 6 TR).

The following event types were used in the deconvolution analysis (as in Chevrier and Schachar, 2010): fixate (F), time-locked to warning-stimuli at the beginning of every trial, left‐ (X) and right-hand (O) response events, time locked to motor responses, successful inhibition (SI), time-locked to the presentation of stop signals, and error detection (Detect) and post-error slowing (PES) events, both time-locked to responses on failed stop trials. Go trials were modeled using (F) and (X) or (O) stimuli. Successful stop trials were modeled using (F) and (SI). Failed stop trials followed by less than median response slowing were modeled with (F), (X) or (O) and (Detect). Failed stop trials followed by greater than median response slowing were modeled with (F), (X) or (O), (Detect) and (PES). Activation maps were estimated by taking the area under the HRF, warped into Talairach space (1mm^3^ resolution), and smoothed using a 6mm full width at half maximum Gaussian kernel.

#### 2.4.1 Assessment of collinearity

Only half of error detection events coincide with post-error slowing, yielding an angle of 60 degrees between these regressors, which is greater than required before multicolinearity becomes a problem (O’Brien, 2007). Variance inflation factor (VIF) is a way of measuring whether regressors that pass collinearity tests with other regressors might instead be collinear with some linear combination of other regressors. VIF is calculated as 1/(1-R^2^), where R is the correlation between the regressor of interest and the most highly collinear combination of other regressors. VIF < 5-10 is considered acceptable in regression analyses. In the current design, the most collinear combination of other event types with our error regressors would be a combination of hand-specific regressors that coincide with error regressors on error trials. However, only one in five responses coincides with error stimuli, and no other regressors in the task could be combined with response regressors to increase their collinearity with error regressors, ensuring a greater angle and therefore less collinearity with error regressors than between error detection and post-error slowing regressors, or VIF < 4/3 (= 1/(1-(cos60)^2^)).

### 2.5 Group level ANOVA analyses

#### 2.5.1 ANOVAs

Single subject activation maps were entered into a random effects ANOVA analysis for ADHD and TD groups. Within group error detection and post-error slowing maps were distributed as t* statistics with 13 degrees of freedom. Group difference maps during error detection and post-error slowing were generated using a nested repeated measures 3-factor ANOVA (group membership, event types, and subjects) to identify significantly different activities between TD and ADHD adolescents. Group difference maps during error detection and post-error slowing were distributed as t* statistics with 26 degrees of freedom.

#### 2.5.2 Correction of multiple comparisons

Output from ANOVA analyses (TD, ADHD, TD-ADHD) were corrected for multiple comparisons using AFNI’s 3dClustSim program (Forman et al., 1995) (spatial correlation estimated from AFNIs corrected 3dFWHMx ‐acf option). This analysis required significant voxels be part of a larger cluster of at least 10.9 original voxels (920 mm^3^) with a minimum Z score of 1.96 for an overall α < 0.05.

The low Z score used for cluster thresholding correction is based on the expected effect sizes from our previous study (Chevrier and Schachar, 2010), which were replicated here with similar statistical power in both TD adolescents and the replication sample of healthy young adults. It has been demonstrated that as threshold is decreased but cluster size is appropriately increased, the rate of false positives remains stable (Slotnick, 2017). Although cluster thresholding has only been validated using Z scores as low as 3.3, we validate the lower Z scores with appropriate cluster sizes used for cluster thresholding here by performing the same analyses and corrections in an independent replication sample of healthy young adults (see **2.8**). If results are false positives due to low power then they will not replicate, whereas if results are true positives then they should replicate even if they are weak. In the ongoing debate about statistical correction approaches, the only point of universal agreement is that there is no stronger or more important basis of validity than replication (Lieberman and Cunningham, 2009; Slotnick, 2017).

### 2.6 Identification of seed and target locations

Seed points for correlations with dorsal striatum in both groups were determined by peak deactivations in dorsal striatum from the TD error detection map. Seed locations for correlation with ventral striatum, SN and raphe nucleus were determined by their peak responses on the TD post-error slowing map. SN activations or correlations within 3 mm (half blurring radius) of subthalamic nucleus (STN), red nucleus or parahippocampus, adjacent to SN, are labeled accordingly in our Tables of results. Seed locations for LC and medial septal nuclei were determined by peak locations of significant negative differences on the TD-ADHD post-error slowing group difference map.

LC seed and target locations (Keren et al., 2009; Tona et al., 2017) were selected based on peak statistics in corrected clusters directly on, or within 1mm of LC atlas locations, or directly between left and right LC. Medial septal nuclei are a visually obvious node superior to the anterior commissure between left and right lateral ventricles (Butler et al., 2014). Results in medial septal nuclei were visually confirmed to be comparable in location to medial septal activations from previous imaging studies (Diaconescu et al., 2017; Iglesias et al., 2013; Shaun Ho et al., 2014).

Raphe nucleus mostly consists of the dorsal nucleus, which runs 6mm superior to the isthmus, and the median nucleus, which runs 10mm inferior to the isthmus, both along the posterior edge of the brainstem (Beliveau et al., 2015; Kranz et al., 2012). Our raphe nucleus seed location was visually confirmed to be comparable in location to raphe nucleus activations from previous imaging studies (Beliveau et al., 2015; Kranz et al., 2012). Due to the lack of a standardized atlas for raphe nucleus and its proximity to other brainstem nuclei, we performed a confirmatory whole-brain correlation analysis (see **2.7.3**), as suggested in (Beissner, 2015). We compared post-error slowing correlations with raphe nucleus in TD with correlations from a previous whole brain connectivity study with raphe nucleus at rest using 3T MRI and validated with PET of 5HT transporter binding (Beliveau et al., 2015), which are also consistent with resting state connectivity measured at 7T (Bianciardi et al., 2016).

The rationale for using activity-based rather than predefined ROIs is that activity is not expected to reflect uniform function across the entire structure of interest, particularly along boundaries to neighboring structures or noise. By contrast, using activity based ROIs as done here runs the risk of double-dipping, which can inflate false positive rates by using regions with already increased signal variance (Kriegeskorte et al., 2009). One approach for validating activity based ROIs for correlation analysis is to use leave-one-out procedures, which estimate the likelihood of replicating the same results. The current replication sample (see **2.8**) offers a much higher standard of validity.

### 2.7 Correlation analyses

#### 2.7.1 Rationale for event related intersubject correlation approach

We used low field MRI (1.5T) and large voxels (3.75 × 3.75 × 6mm), which has a high contrast to noise ratio for estimating within trial activities, but not a high temporal signal to noise ratio required for optimal time-series analyses capable of estimating within-subject inter-regional connectivity (Welvaert and Rosseel, 2013). Although high field strength and time series correlation analyses are the ideal approach for measuring connectivity with brainstem nuclei, the current approach relies on the event related design for separating error detection from post-error slowing and for the identification of seeds for correlation. In-house pilot comparisons using a finger-tapping block design showed that although our 3T system had improved temporal signal to noise, it only had approximately half the contrast to noise ratio for measuring task-related activity at the depth of the thalamus across multiple head coil and pulse sequence combinations. This was due to the normalization of the contrast to noise with respect to baseline appropriate for the current approach (Welvaert and Rosseel, 2013), which increased more than task related signal. Therefore, the 1.5T system used here actually had twice the CNR required for detecting event related BOLD responses compared to the 3T system.

The current low field and large voxel approach affords a very simple analysis, is sufficiently free of distortion in the nuclei of interest (Jezzard and Balaban, 1995; Napadow et al., 2006), sufficiently insensitive to motion and physiological noise (Power et al., 2015), and can detect activity in nuclei as small as a few mm (Chen and Ogawa, 1999).

#### 2.7.2 Correlations

For each group, seed activities were correlated with whole brain activity in their respective maps (*i.e.* seed activities during error detection were correlated with the error detection map; seed activities during post-error slowing were correlated with the post-error slowing map) and with response-phase maps. Statistical parametric maps of B1 (slope) estimates from correlation analyses (distributed as t* statistics with 12 degrees of freedom) were inspected for significant peaks in relevant target regions after correction of multiple comparisons described in **2.5.2**.

#### 2.7.3 Confirmatory correlation analyses

In order to confirm that seed activities in neurotransmitter nuclei identified here reflect associated neurotransmitter function and not noise, we performed confirmatory whole-brain correlation analyses on post-error slowing seed activity in SN and raphe nucleus, in which whole-brain connectivity patterns are well characterized (Beliveau et al., 2015; Bianciardi et al., 2016). Post-error slowing activity in the medial septal seed was inspected for correlation with cholinergic basal forebrain as a way of estimating whether medial septal activity reflected cholinergic function.

### 2.8 Replication sample

We performed the same analyses outlined above in a replication sample of 14 healthy young adults (8 male, mean (±SD) age = 24.0 ±2.8) performing the same task, using the same imaging system and parameters as in the main study. Data was from the placebo condition of a double-blind methylphenidate study, approved by the research ethics board at The Hospital for Sick Children. Subjects gave informed written consent. Identification of seed points and correlation analyses were the same as those described for TD adolescents in the main study except in LC and medial septal nuclei. Correlations with LC used the same seed point as in the main study, as neither TD nor healthy young adults exhibited net deactivation of LC. By contrast, the seed location for medial septal nuclei was identified by peak deactivation in the unthresholded error map based on observations in TD who exhibited net deactivation of medial septal nuclei. Statistical parametric maps of B1 (slope) estimates from correlation analyses were inspected for the presence of whole brain corrected peak correlations in the same structures and in the same direction (positive/negative) as those found in TD adolescents in the current study.

## 3 Results

The 14 TD (mean (±SD) age 15.4 ±1.6) and 14 ADHD (age 13.7 ±2.1) adolescents showed significant age difference (1.71 years, p = 0.024). TD and ADHD groups showed no significant difference in post-error slowing (TD 23.9 ±35.5 ms; ADHD 10.9 ±31.2 ms; p = 0.31), go reaction time (TD 566 ±116 ms; ADHD 663.2 ±155 ms; p=0.072), percent correct go-responses (TD 97.86 ± 2.98%, ADHD 96.79 ± 4.74%, p=0.47) or the percent of successful stop trials (TD 51.41 ± 2.68%, ADHD 52.41 ± 3.66%, p=0.41). The only behavioral difference was in stop signal reaction time, which was longer (35.5 ms, p = 0.039) in the ADHD (233 ±51.0 ms) than in the TD group (198 ±33.0 ms).

fMRI data are available online in the following Mendeley Data public repository: https://data.mendeley.com/datasets/ysc9hfxndp/draft?a=22fbb6fe-803f-44ac-8990-08efd9597e4b

### 3.1 Error detection and post-error slowing activities

Significant activities and group differences at seed locations for correlation analyses are listed in Table 1 (portrayed in S1 Fig.). In TD adolescents, error detection deactivated dorsal striatum and SN, and post-error slowing deactivated ventral pallidum and activated SN, consistent with previous findings in healthy adults (Chevrier and Schachar, 2010). ADHD adolescents activated SN and right amygdala on error detection, and activated ventral pallidum and left amygdala on post-error slowing. The peak group difference in ventral pallidum also overlapped the hypothalamus, medially adjacent to ventral pallidum. As the hypothalamus exerts a dominant influence on autonomic function and on the dopaminergic regulation of SNS function (Menegas et al., 2015), it was included as a potential correlation target and inspected for whole brain corrected correlations with seed activities in neurotransmitter nuclei.

Post-error slowing was also associated with deactivation of raphe nucleus in TD but not ADHD. Consistent with the known level of distortion of the location of raphe nucleus at 1.5T (Jezzard and Balaban, 1995; Napadow et al., 2006), our raphe nucleus ROI was a few mm anterior to known anatomy. We found greater LC and medial septal activity in ADHD compared to TD adolescents. No significant activities or group differences were present in basal forebrain.

**Table 1.** Seed activities during error detection and post-error slowing Z-scores in bold type passed cluster thresholding correction; Location (Talairach space) and magnitude of activation during error detection and post-error slowing for TD, ADHD and group difference (Diff. = TD-ADHD); hTh = hypothalamus, LC = locus coeruleus, pHPC = parahippocampus, SN = substantia nigra, vPallidum = ventral pallidum. See also S1 Fig.

| Target nucleus | Location | Error Detection (Z) |  |  | Post-error slowing (Z) |  |  |
| --- | --- | --- | --- | --- | --- | --- | --- |
|  |  | TD | ADHD | Diff. | TD | ADHD | Diff. |
| Dorsal striatum | 24 -1 1 | **-2.39** | -0.17 | -1.55 | 0.70 | 0.84 | 0.14 |
| vPallidum/hTh | -4 -6 -5 | **3.00** | 0.68 | 1.74 | **-3.31** | 0.76 | **-2.48** |
| SN/pHPC | 12 -13 -13 | **-2.49** | 0.30 | -1.69 | **2.66** | 1.19 | 0.42 |
| Raphe nucleus | -5 -22 -17 | 0.52 | 1.20 | -0.53 | **-2.31** | -0.96 | 1.94 |
| LC | -3 -35 -16 | 1.35 | -0.25 | 1.02 | -1.54 | 1.90 | **-2.41** |
| Medial septal | -3 0 2 | -0.54 | -1.01 | 0.44 | -1.36 | **2.16** | **-2.55** |

### 3.2 Correlations with dorsal and ventral striatum

Consistent with interrupting ongoing processing when errors are detected, deactivation of dorsal striatum on error detection correlated with response-phase activity in right MFG and parietal regions in TD and with MFG in ADHD (Table 2A; S2 Fig. A).

In TD, consistent with a thresholding influence on dorsal striatum, deactivation of ventral pallidum on post-error slowing correlated with greater activation of SN and dorsal pallidum. Consistent with amygdala activity displacing thresholding function in ADHD, activation of ventral pallidum correlated with activation of left amygdala but not dorsal striatum, and was positively instead of negatively correlated with SN (Table 2B; S2 Fig. B).

**Table 2.** Interrupting processing on error detection and thresholding on post-error slowing P-values in bold type passed cluster thresholding correction. DS = dorsal striatum, MFG = middle frontal gyrus, IPL = inferior parietal lobule, pHPC = parahippocampus, dPallidum = dorsal pallidum, R=right ; L=left; Location (Talairach space), p-values and correlation coefficients (r) for TD and ADHD, and p-values for group difference (TD-ADHD=diff). See also S2 Fig.

| Target nucleus | Location | TD p | TD r | ADHD p | ADHD r | p diff |
| --- | --- | --- | --- | --- | --- | --- |
| A Dorsal striatum on error detection, correlations with response phase |  |  |  |  |  |  |
| R MFG | 41 27 40 | **0.0092** | -0.67 | 0.61 | -0.15 | 0.12 |
| R MFG | 46 12 32 | 0.27 | -0.32 | **0.0055** | -0.70 | 0.21 |
| R IPL | 55 -41 46 | **0.00053** | -0.80 | 0.71 | 0.11 | **0.0023** |
| B Correlations with ventral pallidum during post-error slowing |  |  |  |  |  |  |
| SN/pHPC | 17 -17 -7 | **0.0053** | -0.7 | **0.014** | 0.64 | **0.0001** |
| R dPallidum | 18 1 1 | **0.0013** | -0.77 | 0.24 | 0.34 | **0.0013** |
| L Amygdala | -14 -9 -12 | 0.30 | -0.30 | **0.026** | 0.59 | **0.020** |

### 3.3 Correlations with neurotransmitter nuclei

We performed inter-subject correlation analyses on seed activities in SN, raphe nucleus, LC and medial septal nuclei (listed in Table 1) to identify similarities and differences between TD and ADHD groups.

### 3.4 Substantia nigra (SN)

Deactivation of SN on error detection was correlated with greater response-phase activity in raphe nucleus, LC and bilateral hypothalamus in TD (Table 3A; S3 Fig. A). In ADHD, SN activity on error detection was strongly correlated with response phase activity in the amygdala (Table 3A; S3 Fig. A). During error detection, deactivation of SN correlated with deactivation of LC in TD, whereas SN activity in ADHD was correlated with raphe nucleus and hypothalamus, and negatively correlated with medial septal nuclei (Table 3B; S3 Fig. B). During post-error slowing, activation of SN correlated with greater response-phase activity in LC, amygdala and hypothalamus in ADHD (Table 3C; S3 Fig. C). Confirmatory whole brain correlation with post-error slowing activity in SN (portrayed in S6 Fig.) showed diffuse bilateral correlations with limbic, striatal and neocortical regions in TD, whereas striatal correlations with SN were absent in ADHD.

**Table 3.** Correlations with substantia nigra (SN) Location (Talairach space), for TD (TD p) and ADHD (ADHD p), correlation (r), and p-values for group difference (diff=TD-ADHD). P-values in bold type passed cluster thresholding correction. R=right: L=left; hTh = hypothalamus; LC = locus coeruleus; SN = substantia nigra. See also S3 Fig..

fMRI of errors in ADHD
| Target nuclei | Location | TD p | TD r | ADHD p | ADHD r | p diff |
| --- | --- | --- | --- | --- | --- | --- |
| A SN on error detection, correlations with response-phase |  |  |  |  |  |  |
| L Amygdala | -21 -7 -21 | 0.63 | 0.14 | **1.8e-6** | 0.93 | **0.0004** |
| Raphe nucleus | -4 -19 -13 | **0.0018** | -0.76 | 0.78 | -0.084 | **0.032** |
| LC | -3 -35 -16 | **0.029** | -0.58 | 0.22 | 0.35 | **0.016** |
| Bi hTh | 0 -2 -9 | **0.003** | -0.73 | 0.26 | -0.32 | 0.16 |
| B Correlations with SN during error detection |  |  |  |  |  |  |
| Raphe nucleus | -1 -24 -15 | 0.57 | 0.16 | **0.0092** | 0.67 | 0.13 |
| LC | 6 -37 -24 | **0.01** | 0.66 | 0.53 | 0.65 | 0.97 |
| Medial septal | -2 8 5 | 0.33 | 0.28 | **0.019** | -0.62 | **0.018** |
| R hTh | 5 -1 -13 | 0.38 | -0.26 | **0.00033** | 0.82 | **0.0008** |
| C SN on post-error slowing, correlations with response-phase |  |  |  |  |  |  |
| L Amygdala | -25 -3 -18 | 0.035 | 0.54 | **0.0056** | 0.70 | 0.54 |
| R LC | 7 -37 -26 | 0.43 | -0.17 | **0.049** | 0.53 | 0.074 |
| L hTh | -3 -7 -6 | 0.91 | -0.034 | **0.046** | 0.54 | **0.03** |

### 3.5 Raphe nucleus

In both groups, deactivation of raphe nucleus during post-error slowing correlated with less amygdala activity during post-error slowing (Table 4A; S4 Fig. A) and more amygdala activity during response-phases (Table 4B; S4 Fig. B). Deactivation of raphe nucleus during post-error slowing correlated with greater activation of SN in TD (Table 4A; S4 Fig. A). Greater deactivation of raphe nucleus during post-err(S6 Fig.)or slowing also correlated with less activation of SN and greater activation of medial septal nuclei and hypothalamus during response-phases in ADHD, but with less response-phase activity in hypothalamus in TD (Table 4B; S4 Fig. B).

**Table 4.** Correlations with raphe nucleus Location (Talairach space), p-values and correlation coefficients (r) for TD and ADHD, and pvalues for group difference (TD-ADHD=diff). P-values in bold type passed cluster thresholding correction. R=right; L=left; Bi=bilateral; hTh = hypothalamus; pHPC = parahippocampus; RedNuc = red nucleus; SN = substantia nigra. See also S4 Fig.

| Target nuclei | Location | TD p | TD r | ADHD p | ADHD r | p diff |
| --- | --- | --- | --- | --- | --- | --- |
| A Correlations with raphe nucleus during post-error slowing |  |  |  |  |  |  |
| L Amygdala | -21 -7 -18 | **0.00013** | 0.85 | 0.06 | 0.51 | 0.10 |
| L Amygdala | -24 -4 -17 | **0.0026** | 0.74 | **5.4e-5** | 0.87 | 0.37 |
| R Amygdala | 20 0 -20 | **0.00065** | 0.80 | 0.28 | 0.31 | 0.069 |
| R Amygdala | 21 -2 -12 | 0.97 | 0.0098 | **0.00022** | 0.83 | **0.0058** |
| L SN/RedNuc | -7 -19 -10 | 0.73 | 0.10 | **0.022** | -0.60 | 0.10 |
| R SN | 12 -14 -9 | **0.035** | -0.57 | 0.88 | -0.043 | 0.16 |
| R SN/pHPC | 12 -20 -10 | 0.89 | -0.041 | **0.0015** | -0.76 | **0.025** |
| B Raphe nucleus during post-error slowing, correlations with response-phase |  |  |  |  |  |  |
| L Amygdala | -17 -5 -15 | **0.00046** | -0.81 | 0.88 | 0.044 | **0.006** |
| L Amygdala | -22 -8 -17 | **0.024** | -0.60 | **0.00035** | -0.82 | 0.28 |
| R Amygdala | 30 -2 -18 | **8.0e-5** | -0.86 | 0.47 | -0.21 | **0.011** |
| R SN | 9 -14 -10 | 0.16 | -0.39 | **0.0045** | 0.71 | **0.0023** |
| Medial septal | 7 6 4 | 0.41 | -0.24 | **0.018** | -0.62 | 0.26 |
| L hTh | -5 -6 -10 | **0.0065** | -0.69 | 0.055 | 0.52 | **0.0008** |
| Bi hTh | 5 -2 -12 | 0.52 | -0.19 | **0.0045** | 0.71 | **0.011** |

In TD, confirmatory whole brain correlation with post-error slowing activity in raphe nucleus replicated previous findings using PET and resting state fMRI (Beliveau et al., 2015; Bianciardi et al., 2016) of positive correlations in parahippocampus, anterior insula, ACC/OFC, superior temporal and precuneus, and negative correlations in bilateral pre/post/paracentral and left superior frontal gyrus (S7 Fig.). The only correlation from Beliveau et al (2015) that was not present in TD was with right superior parietal lobule. Raphe nucleus activity in the replication sample exhibited the same whole-brain correlations. Correlations with raphe nucleus were diminished in ADHD except in the amygdala and parahippocampus, and were significantly heightened in the anterior insula.

### 3.6 Locus coeruleus (LC)

During post-error slowing, deactivation of LC correlated with hypothalamus and raphe nucleus in TD (Table 5A; S4 Fig. C). In ADHD, activation of LC on post-error slowing was correlated with less activation of right amygdala and greater activation of the inhibitory control network involving right inferior frontal gyrus and caudate nucleus (Aron et al., 2004; Chevrier et al., 2007) (Table 5A; S4 Fig. C). Greater activation of LC on post-error slowing in ADHD was also correlated with greater response-phase activity in SN, raphe nucleus and the amygdala (Table 5B; S4 Fig. D).

**Table 5.** Correlations with locus coeruleus (LC) Location (Talairach space), p-values and correlation coefficients (r) for TD and ADHD, and pvalues for group difference (TD-ADHD=diff). P-values in bold type passed cluster thresholding correction. R=right; L=left; IFG = inferior frontal gyrus; hTh = hypothalamus; SN = substantia nigra. See also S4 Fig.

| Target nuclei | Location | TD p | TD r | ADHD p | ADHD r | p diff |
| --- | --- | --- | --- | --- | --- | --- |
| A Correlations with locus coeruleus during post-error slowing |  |  |  |  |  |  |
| R Amygdala | 20 -8 -13 | 0.86 | -0.05 | **0.014** | -0.64 | 0.097 |
| Raphe | 1 -26 -18 | **0.0053** | 0.70 | 0.14 | 0.42 | 0.33 |
| L hTh | -3 -2 -10 | **0.0011** | 0.78 | 0.75 | -0.09 | **0.0078** |
| R IFG 45 | 58 20 15 | 0.95 | 0.02 | **0.0042** | 0.73 | **0.033** |
| R Caudate | 19 17 11 | 0.64 | 0.14 | **0.00026** | 0.28 | 0.74 |
| B Locus coeruleus during post-error slowing, correlations with response-phase |  |  |  |  |  |  |
| L Amygdala | -22 -10 -9 | 0.61 | 0.15 | **0.00078** | -0.65 | **0.03** |
| R SN | -7 -13 -10 | 0.074 | -0.49 | **0.015** | -0.63 | 0.63 |
| Raphe | 2 -24 -16 | 0.15 | -0.15 | **0.022** | -0.74 | 0.061 |

### 3.7 Medial septal nuclei

In TD, deactivation of medial septal nuclei correlated with greater amygdala activity during post-error slowing (Table 6A; S4 Fig. E) and less amygdala and LC activity during response-phases (Table 6B; S4 Fig. F). In ADHD, post-error slowing activation of medial septal nuclei correlated with SN activity during post-error slowing (Table 6A; S4 Fig. E) and less amygdala but greater hypothalamus activity during response-phases (Table 6B; S4 Fig. F). Medial septal activity was correlated with bilateral basal forebrain in ADHD, consistent with medial septal activity reflecting an influence of the cholinergic system (Table 6A; S4 Fig. E).

**Table 6.**
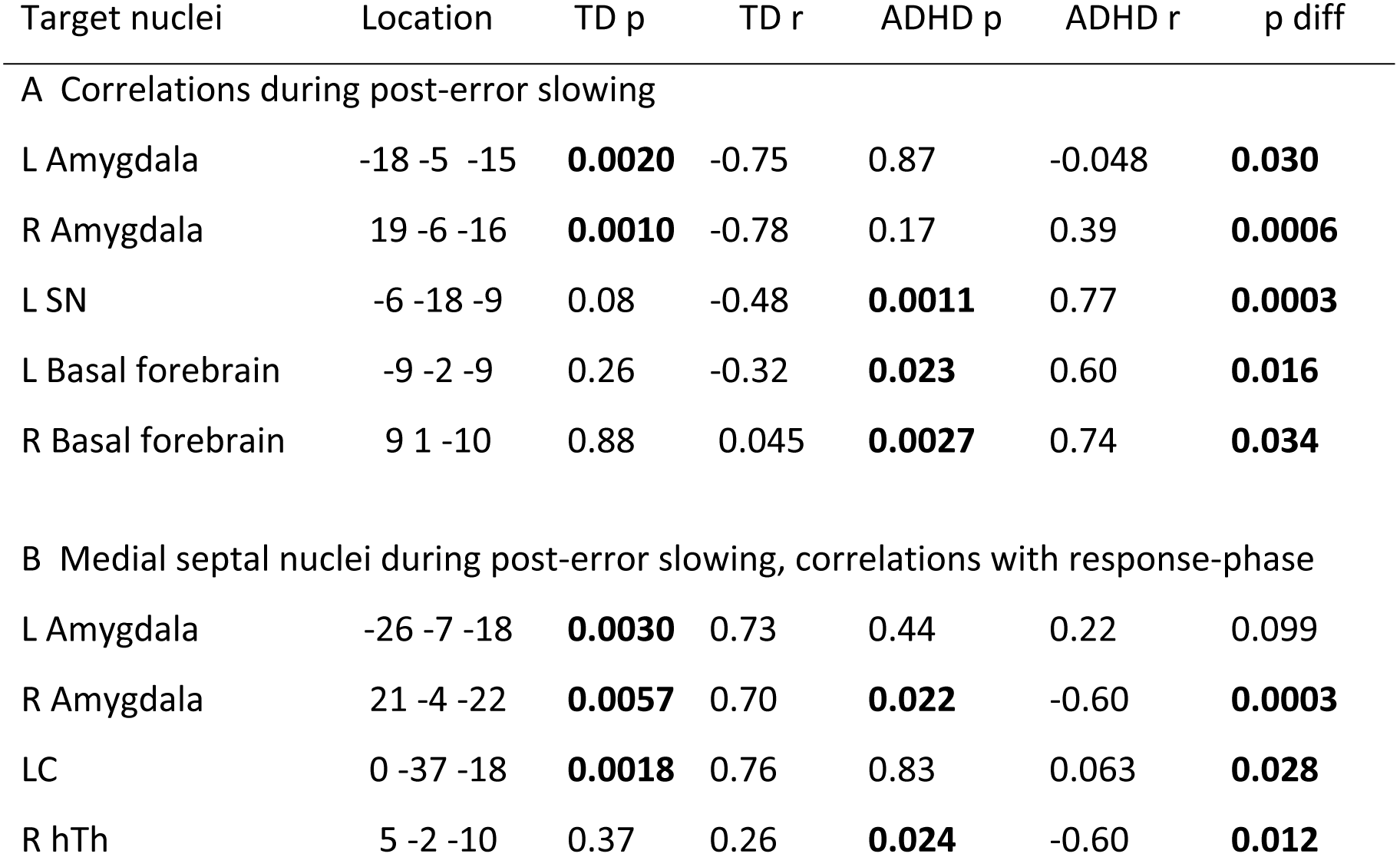
Correlations with medial septal nuclei Location (Talairach space), p-values and correlation coefficients (r) for TD and ADHD, and pvalues for group difference (TD-ADHD=diff). P-values in bold type passed whole brain correction. * = values from separate ROI analysis with basal forebrain; R=right; L=left; hTh = hypothalamus; LC = locus coeruleus; SN = substantia nigra. See also S4 Fig.

### 3.8 Replication of TD correlations in healthy adults

Behavioral performance in the replication sample was in normal range (RT = 538.5 *±* 92.0 ms; SSRT = 218.3 *±* 38.2 ms; post-error slowing = 12.8 ± 38.6 ms; percent successful stop trials = 50.18 *±* 1.34%; percent correct go response = 98.47 *±* 1.57%). Table 7 lists the same pattern of correlations in the replication sample (S5 Fig.) as in TD adolescents from the main study. Three correlations did not survive whole brain correction (ventral pallidum with SN during post-error slowing, raphe nucleus during post-error slowing with right amygdala during response-phases, and medial septal nuclei with left amygdala during post-error slowing), but were in the same direction (positive/negative) as in the TD group. Correlations with ventral pallidum and one other correlation (raphe nucleus during post-error slowing with right amygdala during response-phases) were not related to the slope, but rather the baseline term, indicating that these correlations succeeded in capturing the expected relative levels of mean group activity in seed and target regions, but failed to capture the trend of inter-subject variability. One correlation was present in the replication sample but not in the TD group (raphe nucleus with LC during post-error slowing), and three bilateral amygdala correlations in TD (raphe during post-error slowing with response phase, and medial septal nuclei during post-error slowing with post-error slowing and response-phase) were only present in one hemisphere in the replication sample. Nonetheless, the replication sample contained all the same correlations in the same direction (positive/negative) as in TD.

**Table 7.**
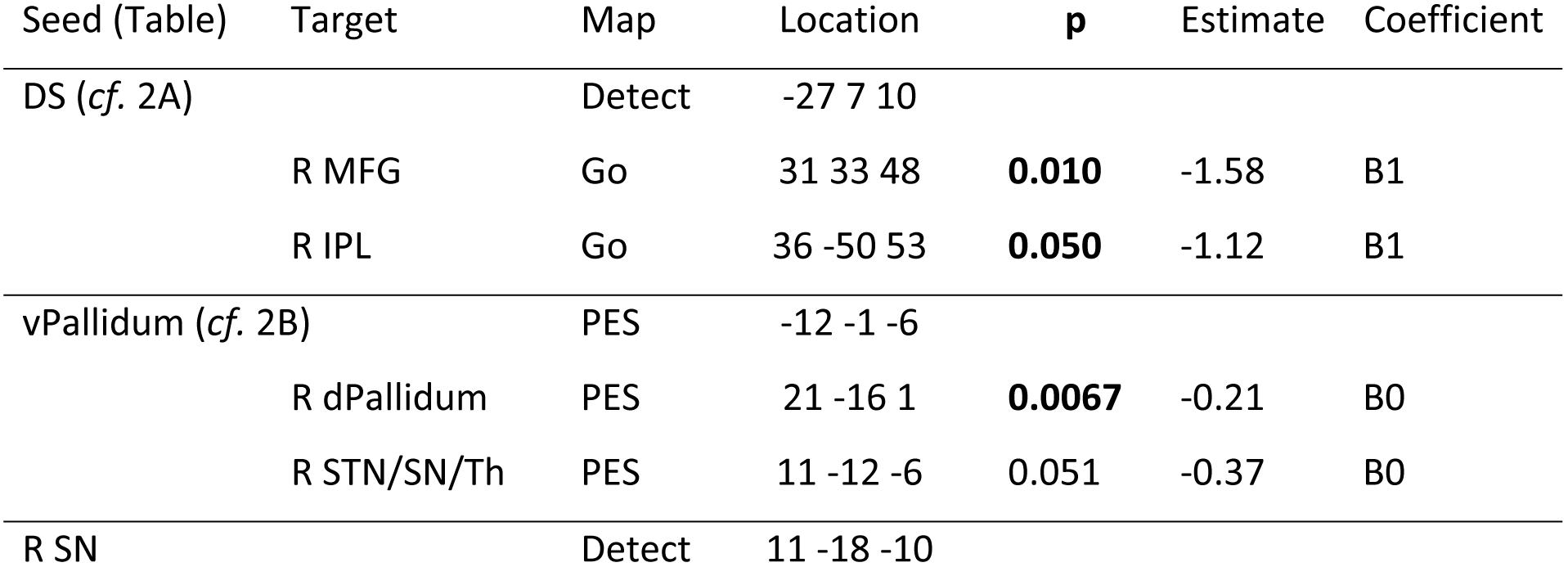
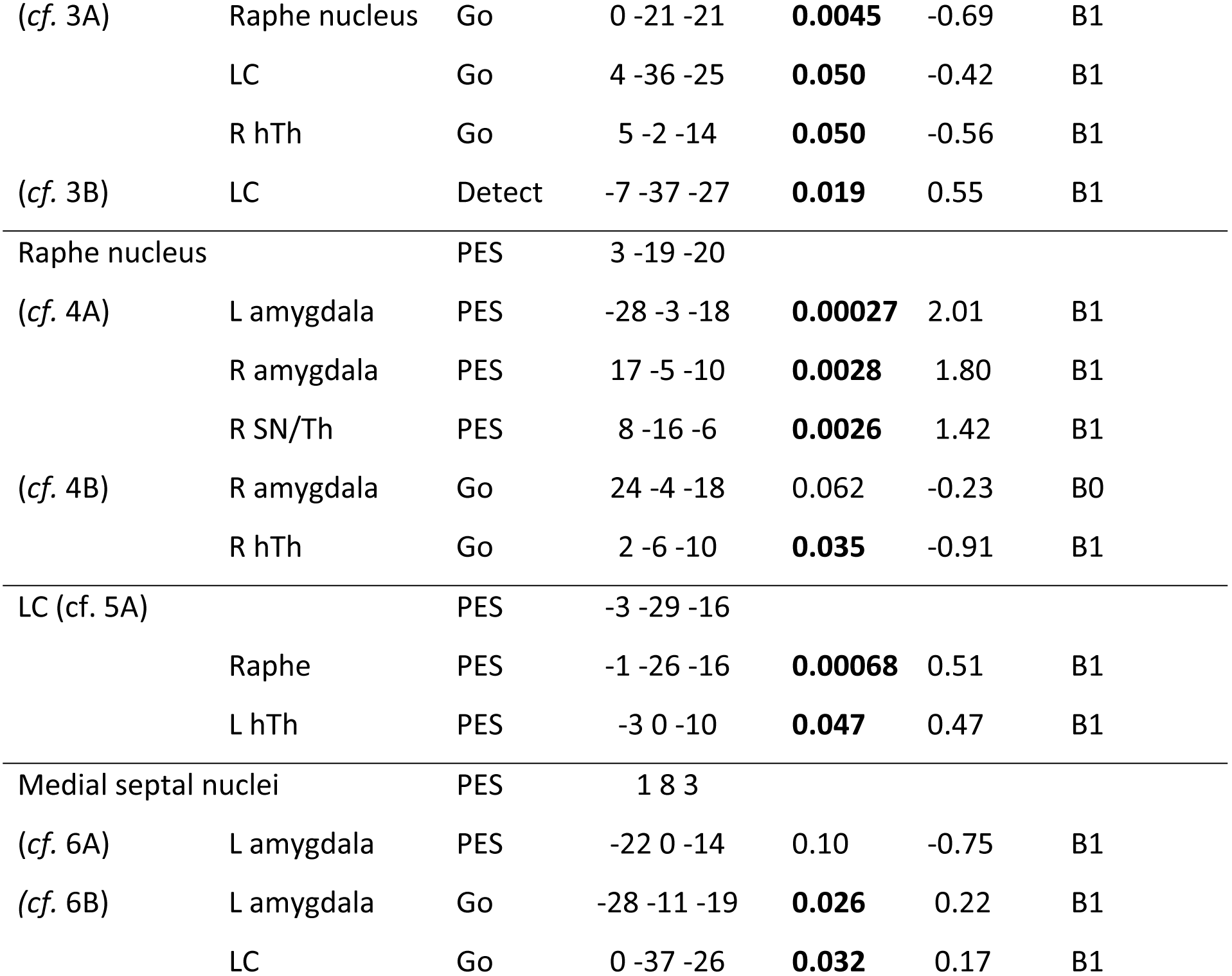
Replication of correlations from TD in healthy adults. All correlations in TD adolescents (Tables 2-6) were also present in the same directions (positive/negative) in the replication sample. Table shows seed and target regions (*cf.* tables with corresponding correlations in TD), locations (Talairach space), phase of seed and target activities for correlation (Go = response-phase, Detect = error detection, PES = post-error slowing), p-values and coefficient estimates (small volume corrected, r=3mm), and whether correlation was with baseline (B0) or slope (B1) coefficient. P-values in bold type passed cluster thresholding correction. R=right; L-left; DS=dorsal striatum; dPallidum=dorsal pallidum; hTh = hypothalamus; IPL=inferior parietal lobule; LC = locus coeruleus; MFG=middle frontal gyrus; SN=substantia nigra; STN=subthalamic nucleus; Th = thalamus; vPallidum=ventral pallidum. See also S5 Fig.

## 4 Discussion

This is the first study to separate error detection and post-error slowing in TD and ADHD adolescents, and to identify the presence or absence of striatal thresholding influences at the core of reinforcement learning function. Results in TD replicated our previous findings of deactivations in dorsal and ventral striatum during error detection and post-error slowing. Correlation analyses in TD were consistent with dorsal striatum interrupting ongoing processing when errors are detected, and with ventral striatum affecting thresholds in dorsal striatum and SN during post-error slowing. In ADHD, we found heightened amygdala activity on errors similar to that which has been observed in tasks using more emotional stimuli (Maier et al., 2014), and correlation analyses were consistent with this activity preventing thresholding on post-error slowing. We also found heightened activities and altered inter correlations among multiple neurotransmitter nuclei consistent with an altered competition for control of DA in ADHD. Identifying reinforcement learning influences in the striatum and among competing neurotransmitter nuclei during prediction errors is crucial to understanding normal and altered development because the integrated function of these systems strongly influence how all functional networks are fine-tuned based on experience.

### 4.1 Methodological considerations

The current approach employed intersubject correlation analyses at low field using large voxels, and low statistical thresholds for estimating connectivity in small nuclei that are normally measured with time series approaches at high field with smaller voxels and more stringent statistical thresholds. Given these differences, it is important to contextualize the interpretation of results in terms of the rationale for the methods we used, and its similarities and differences with previous approaches.

The low threshold and large cluster size correction used here were based on the expected effect sizes from our previous study (Chevrier and Schachar, 2010). We found considerable intersubject variability of activity in the nuclei studied here, and strong inter-correlations combined with successful replication indicate that this variability in activity reflects variability of function and not simply measurement error. Therefore, the low threshold and large cluster size required for identification of seed points for correlation are appropriate and reproducible, and will not likely be much improved by increasing the number of subjects. Further, despite significant group differences, there was considerable overlap between the levels of TD and ADHD activity in the nuclei studied here, which could indicate overlapping function. However, different intersubject correlations indicate that comparable levels of activity do not reflect comparable functional influences in both groups.

Despite an absence of measurable group differences in the magnitude of post-error slowing, which was not predicted in the relatively small sample size used here, TD and ADHD groups exhibited highly distinct patterns of activity and intersubject correlations on errors followed by greater than median post-error response times. It is possible that the current approach simply does not appropriately capture error magnitude in the SST. Indeed, many trial by trial effects could have contaminated our rudimentary median split of post-error slowing with non-error magnitude related functions (Dupuis et al., 2018). However, models of trial by trial effects (e.g. (Den Ouden et al., 2009; Elliott et al., 2000; O’Doherty et al., 2003; Ullsperger et al., 2014)) are either models of reinforcement learning itself or pertain to modular functions that would be interrupted and adjusted on errors, and are therefore higher order effects on the core neural mechanisms identified here.

Although the current replication of TD correlations in healthy adults validates the use of peak BOLD responses rather than predefined ROIs as seeds for correlation (Kriegeskorte et al., 2009), our results further demonstrate the necessity of this approach for identifying thresholding function on errors. Several neurotransmitter nuclei exhibited diffuse correlations in distributed target regions, such as SN correlations throughout the striatum and neocortex. However, correlations directly involved in interrupting and adjusting task-related functions were far more localized. For example, while SN correlated with nearly the entire striatum on post-error slowing, deactivation of ventral striatum only correlated with pallidal output from the same part of dorsal striatum that deactivated on error detection, which correlated specifically with task-related response phase activity. Predefined anatomy based ROIs would not have captured these effects, which likely reflect activity level dependent bistable firing properties of subcortical gating mechanisms that have been proposed to drive ADHD symptoms (Levy, 2004). The dissociation of diffuse effects of neurotransmitter nuclei from more specific activity level dependent thresholding influences on task-related processing demonstrates the necessity of the event-related activity based ROI approach.

Our results also show that the low field large voxel approach can successfully identify rapidly changing connectivities with specific brainstem nuclei on errors in the SST. In particular, confirmatory whole brain correlation analyses with post-error slowing activity in SN and raphe nuclei were consistent with their known connectivity patterns, providing definitive evidence that these activities reflect the same kind of neurotransmitter function that has been identified with PET and time series analyses using high-field fMRI (Beliveau et al., 2015; Bianciardi et al., 2016), and rules out previous assertions that SN activity identified with this approach instead reflects subthalamic nucleus (Danielmeier and Ullsperger, 2011). LC is large enough to be detectable using low field large voxel fMRI (Chen and Ogawa, 1999) and has similar levels of neighboring noise (Brooks et al., 2013) and less distortion (Jezzard and Balaban, 1995; Napadow et al., 2006) than raphe nucleus across multiple field strengths. Although the lack of established whole brain correlation patterns with LC do not allow for the same validation of function as results in SN and raphe nucleus, the principled nature of LC responses and correlations combined with replication of correlations from TD in healthy adults support these activities reflecting LC function and not neighboring noise. Medial septal nuclei are more than large enough for generating detectable event-related BOLD signal, have no other adjacent nuclei along the anterior-posterior axis of geometric distortion, and neighboring noise is comparable to levels in raphe nucleus across multiple field strengths (Brooks et al., 2013; Hutton et al., 2011). Further, elevated medial septal activity in ADHD correlated with cholinergic basal forebrain during post-error slowing, consistent with these activities reflecting medial septal function.

Given the large number of correlations inherent to this study, each of which imposes a compounding cost on the probability of replication, the replication of all TD correlations with the same nuclei in the same direction (positive/negative) at comparable levels of significance in healthy adults provide definitive evidence that the statistical corrections used here were appropriate and that results in these nuclei reflect signal and not noise. Only one correlation was present in the replication sample but not in TD (raphe nucleus with LC during post-error slowing), although subthreshold correlation was positive in TD as in the replication sample. This level of replication of correlations from TD in healthy adults who were on average 8.6 years older, also rules out concerns that group differences compared to ADHD could be the result of a 1.7 year mean age difference, or other idiosyncratic subject matching issues. On the contrary, intersubject variability is an important form of contrast in the intersubject correlation approach, enhancing rather than diminishing important functional group differences. The only remaining alternative interpretation of the current replication would be that these results simply reflect noise that rose above the low threshold used here, which can be ruled out based on the distinct pattern of results in ADHD. If the level of replication we observed in TD were simply noise, then the same results would also be replicated to a similar degree in ADHD.

### 4.2 Thresholding on post-error slowing in TD, not ADHD

We initially hypothesized that the same part of dorsal striatum involved in magnitude-related thresholding must first interrupt ongoing processing when errors are detected. In TD, we found that deactivation of dorsal striatum on error detection was indeed correlated with response phase activity in right frontoparietal regions that are known to activate during response phases in the SST. Deactivation of dorsal striatum on error detection was not significant in ADHD, and was correlated with frontal but not parietal response-phase activity. The presence of prefrontal but not parietal response-phase activity is consistent with decreased parietal involvement in task-related processing that has been observed in ADHD (Christakou et al., 2013; Tomasi and Volkow, 2012). Despite dorsal striatum interrupting response phase processing on error detection in both groups, post-error slowing activities triggered by error detection were highly distinct in TD and ADHD.

Correlation of ventral pallidum with SN and dorsal striatum during post-error slowing in TD is consistent with striatal thresholding influences necessary for reinforcement learning (Horvitz, 2002). Opposite activity in these structures during error detection would explain why thresholding effects have not been apparent from studies of resting state connectivity or task based approaches that have not separated error detection from post-error adjustment.

In ADHD, correlation of ventral striatum with the amygdala and not with dorsal striatum supports our initial hypothesis that heightened amygdala activity drives limbic-motor interfacing that would prevent thresholding. Although it is possible that post-error slowing simply does not capture error magnitude in ADHD as in TD adolescents, the current results are consistent with opposite ventral striatal activity that has been found in studies that have experimentally controlled prediction error magnitude (Furukawa et al., 2014; Wilbertz et al., 2012).

The ventral pallidal seed we used for correlation, identified by peak group difference activity during post-error slowing, also overlapped the hypothalamus, medial to ventral pallidum. The hypothalamus is involved in autonomic function, and is a major source of input to dopaminergic projections to the SNS (Menegas et al., 2015). Gating influences from the SNS can be strongly influenced by DA-amygdala interactions, and determine the scope of modular functions that can participate in globally integrated processing at any given time(Fudge and Emiliano, 2003; Haber et al., 2000; Morrison et al., 2017). Altered activities and correlations with the hypothalamus, combined with evidence of limbic-motor interfacing in place of error magnitude related thresholding, would support very different forms of globally integrated processing in ADHD compared to TD

### 4.3 Altered competition for control of DA in ADHD

#### 4.3.1 Altered activities and correlations with SN and LC

Group differences in post-error slowing activities and correlations with SN and LC are consistent with emerging evidence that prediction errors trigger controlled DA-driven adjustment in healthy subjects and more stochastic NE-driven adjustment in ADHD (Aston-Jones and Cohen, 2005; Del Campo et al., 2011; Hauser et al., 2016, 2014). Activities in LC (NE) and medial septal nuclei (Ach) signal surprise (Yu and Dayan, 2003) and drive externally-directed attention and learning about context (Gu, 2002; Kimura, 1999; Kobayashi et al., 2000). Therefore, deactivation of LC and medial septal nuclei in TD would facilitate internally-directed attention for reinforcement learning. In ADHD, activation of LC would instead drive externally-directed attention and learning about context that is at odds with reinforcement learning.

Consistent with LC rather than SN-driven function, activation of LC in ADHD was specifically correlated with right inferior frontal gyrus and caudate, which is precisely the network most implicated in response inhibition (Aron et al., 2004; Chevrier et al., 2007). Therefore, heightened LC activity in ADHD appears to be driving an urgent attempt to stop, after errors have already been committed, which would result in very different reinforcement of function compared to the SN driven reinforcement we observed in TD. Although activity in right inferior frontal gyrus was significantly different between groups, the level of activity in the caudate nucleus, often used as an index of the kind of reinforcement learning function observed in TD, was actually similar in the ADHD group. Despite similar levels of activity, the qualitative differences in SN and LC correlations clearly illustrates that caudate activity during post-error slowing reflects very different functions in TD and ADHD.

It was recently shown that the hippocampus receives DA input from LC rather than from dopaminergic SN (Kempadoo et al., 2016). The division of DA projections from SN to the striatum, necessary for implicit learning, and from LC to the hippocampus, necessary for explicit learning, point to the importance of imaging multiple neurotransmitter nuclei simultaneously on prediction errors. The heightened LC response to post-error slowing we observed in ADHD is consistent with an altered balance of explicit, hippocampus-dependent learning, and implicit striatum-dependent learning (Shohamy et al., 2008). Consistent with this hypothesis, whole-brain correlation of post-error slowing activity in SN showed bilateral limbic, striatal and neocortical correlations consistent with ascending DA pathways in TD, whereas the striatal component of these correlations was almost entirely absent in ADHD. This result is in line with the fact that expected nigrostriatal deficits associated with ADHD have not been attributable to inherent differences in striatal DA receptor density, but are thought to arise from the effects of decreased tonic and increased phasic catecholamine responses on striatal gating function via the SNS (Del Campo et al., 2013).

The relatively stark absence of striatal correlations with SN during post-error slowing in ADHD, combined with heightened responses and altered correlations with the amygdala and non-dopaminergic neurotransmitter nuclei are consistent with gating conditions in the SNS similar to a functional lesioning of nigrostriatal thresholding influences during the brief but crucial time window when reinforcement learning must occur. Animal lesion studies have shown that without appropriate function in the striatum at the moment that errors occur, learning and adjustment do not take place despite apparently functioning perceptual and cognitive systems with full access to the history of events (Atallah et al., 2007). Humans certainly have access to levels of associative learning not available to animals. However, the lack of implicit learning effects that normally coincide with appropriate responses in limbic and neurotransmitter nuclei, which are tightly coupled with and actively monitor our visceral states, would lend a very different sense of reliability and informativeness to the same feedback. Further, lack of DA-driven error-related activity in Parkinson’s disease has been associated with total lack of awareness of errors (Palermo et al., 2017).

Imaging and behavioral data have been rigorously linked to conceptual frameworks of model-based Bayesian inference and model-free reinforcement learning (Den Ouden et al., 2009; Elliott et al., 2000; O’Doherty et al., 2003; Ullsperger et al., 2014). These and other approaches are capable of quantifying departures from history dependent task adjustments (Fischer and Ullsperger, 2017; Hauser et al., 2016). Models of task adjustments naturally assume that behavior and neural activity are directed towards performance on the task itself, which is necessary for establishing a metric of accuracy for fitting models to behavior. Our results are in agreement with these assumptions being valid in TD and healthy young adults. But what if the relevance of neural differences in ADHD were less about quantitative differences in task-directed processing and more about qualitative differences in what that processing reflects? Altered correlations found here, and widely consistent findings of altered connectivity in ADHD (Cao et al., 2014; Choi et al., 2013; Liston et al., 2011; Ma et al., 2016; Murias et al., 2007; Tian et al., 2006; Tomasi and Volkow, 2012), suggest that it might indeed be the latter.

Heightened rather than suppressed activities in LC and medial septal nuclei like we observed in ADHD on post-error slowing are known to direct attention towards the external environment and facilitate learning about context rather than the internal details of any task (Gu, 2002; Kobayashi et al., 2000; Yu and Dayan, 2003). Further, seed activities in all the neurotransmitter nuclei studied here exhibited significant group differences in their correlation with the hypothalamus, which projects to and exerts a strong influence on all of these nuclei (Li et al., 2014). These differences in activity and patterns of correlation support the notion that activities in neurotransmitter nuclei are driving mutually exclusive operating conditions in ADHD compared to TD, which support distinct forms of attention, cognition and behavior. This work can help to delineate the kinds of functions that altered error activity actually reflects in ADHD, which could help identify alternative neurocognitive models and alternative targets and strategies for intervention.

#### 4.3.2 Altered activation and correlations with medial septal nuclei

Heightened medial septal activity on post-error slowing in ADHD was negatively correlated with amygdala activity during post-error slowing and response-phases, suggesting that this activity may help to suppress heightened emotional responses to errors. By contrast, medial septal deactivation showed weakly positive correlation with the amygdala in TD. Deactivation of medial septal nuclei may therefore be an aversive dimension of habit-breaking, negative reinforcement in healthy individuals, as activation of medial septal nuclei is highly pleasurable and habit-forming (Mamad et al., 2015; Olds and Milner, 1954; Partridge et al., 2002). Ach function is altered in ADHD and interacts with other neurotransmitter systems differently than in healthy subjects (Beane and Marrocco, 2004; Johansson et al., 2013; Tharoor et al., 2008; Wallis et al., 2009). Heightened activation of medial septal nuclei and altered correlation with the amygdala in ADHD might indicate an inherent resistance to the aversive and habit-breaking effects of negative reinforcement.

#### 4.3.3 Distinct and similar correlations with raphe nucleus

5HT function in raphe nucleus has joint influences with DA on the rewarding properties of prediction errors (Fischer and Ullsperger, 2017) and competes with NE for control of DA in a way that can maintain effortful DA-regulated behavior (i.e. decrease rest duration between phases of exertion) (Auclair et al., 2004; Meyniel et al., 2016). Raphe nucleus deactivated on post-error slowing in TD but not ADHD, who exhibited significantly different correlations with SN activity, consistent with an altered competition for DA (Auclair et al., 2004).

The only correlations with neurotransmitter nuclei common to both groups were between raphe nucleus and the amygdala. This result suggests that the relationship between effort cost and emotional regulation may be a shared dimension of variability in otherwise highly distinct behavioral management landscapes associated with TD and ADHD.

### 4.4 Conclusions

The striatal and neurotransmitter nuclei identified here must function as an integrated unit (Fischer and Ullsperger, 2017; Horvitz, 2002; Kempadoo et al., 2016), and fMRI offers the ability to image them in an integrated context. Several correlations with neurotransmitter nuclei were more significant than correlations with regions directly involved in task performance or striatal thresholding, indicating the consistent and dominant influence of interacting neurotransmitter nuclei on prediction errors. In ADHD, the cumulative effects of limbic-motor interfacing during the crucial time window that normally fine-tunes behavior based on feedback would contribute to altered developmental trajectories in ADHD. These alterations include preparatory deactivation instead of activation of task-related networks (Bhaijiwala et al., 2014), persistent inhibitory control deficits (Schachar et al., 1995), and lack of potency of behavioral interventions (Wolraich et al., 2011).The ability to simultaneously observe the interplay among multiple neurotransmitter nuclei and striatal gating functions during error detection and post-error slowing can improve our understanding of these systems during normal and altered reinforcement learning, and inform predictions and monitoring of the response of these systems to pharmaceutical and other interventions.

## Acknowledgements

This work was supported by a grant to R. S. from the Canadian Institutes of Health Research (MOP 82796).

---

**S1 Fig.**
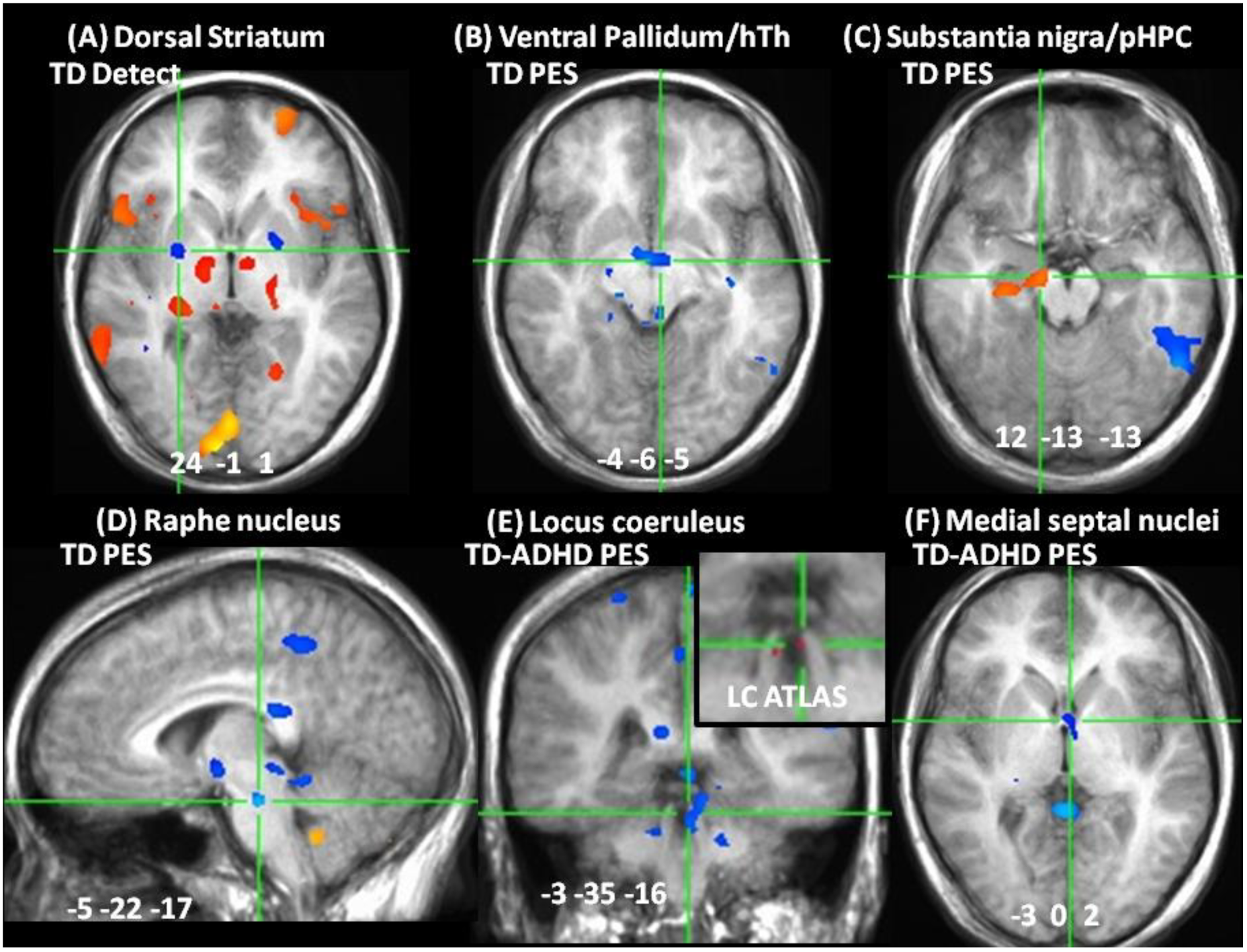
**Seed activities related to Table 1.** (A) Deactivation of dorsal striatum in TD adolescents during error detection (TD Detect). (B) Deactivation of ventral pallidum in TD during post-error slowing (TD PES). (C) Activation of substantia nigra in TD during post-error slowing (TD PES). (D) Deactivation of raphe nucleus in TD adolescents during post-error slowing (TD PES). (E) Negative group difference in locus coeruleus during post-error slowing (TD-ADHD PES; inset shows locus coeruleus (LC) atlas in red). (F) Negative group difference during post-error slowing in medial septal nuclei (TD-ADHD PES). Activation maps portray percent BOLD estimates after whole brain correction (red/yellow = activation, blue = deactivation). Locations in Talairach space, portrayed in radiological space (left = right); hTh = hypothalamus; pHPC = parahippocampus.

**S2 Fig.**
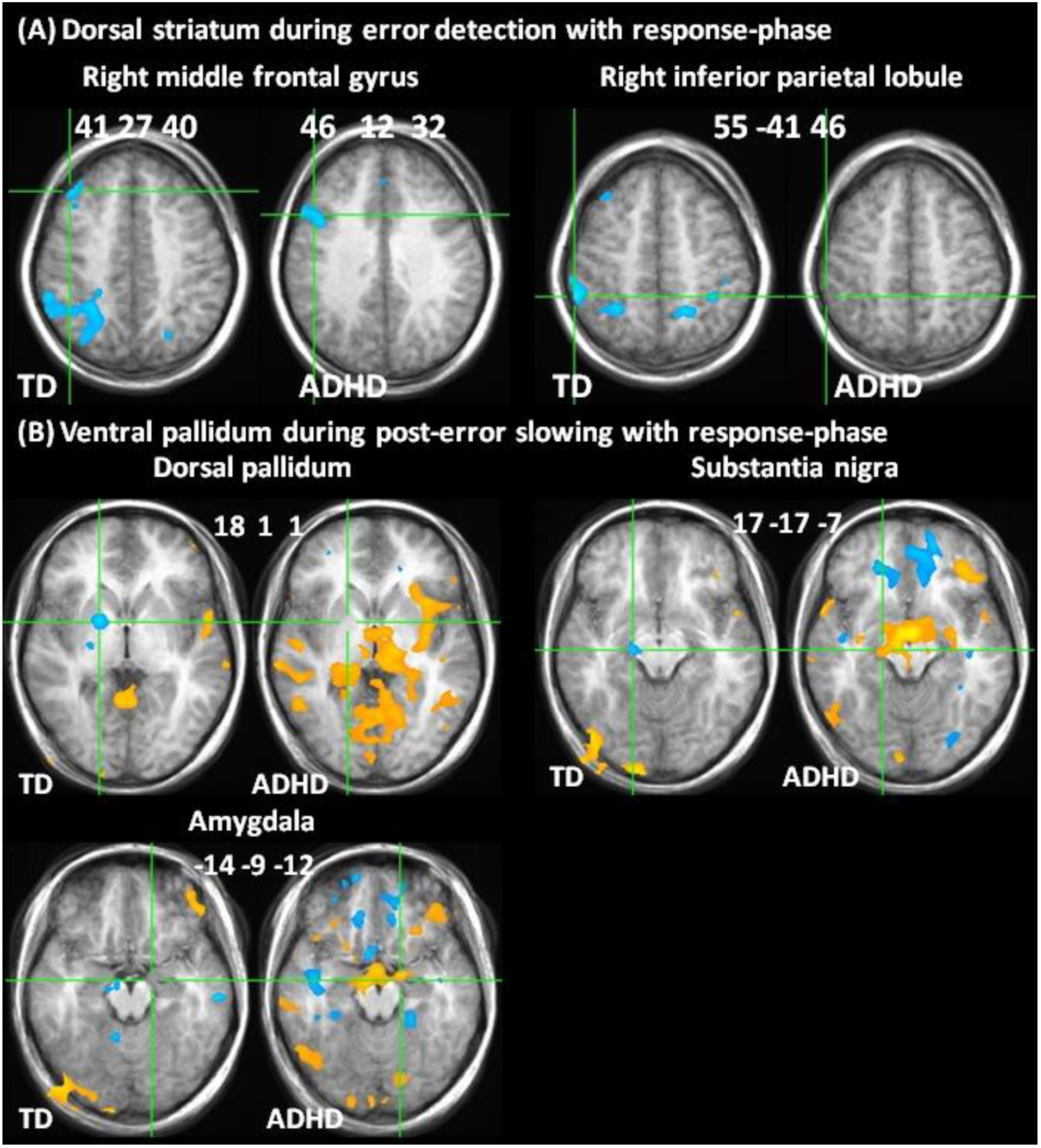
**Correlations with dorsal striatum and ventral pallidum related to Table 2**. (A) Correlations of dorsal striatum during error detection with right middle frontal gyrus and inferior parietal lobule during response-phases. (B) Correlations of ventral pallidum with dorsal pallidum, substantia nigra and amygdala during error detection. Correlation maps portray B1 estimates after whole brain correction (red/yellow = positive, blue = negative correlation). Locations in Talairach space, portrayed in radiological space (left = right).

**S3 Fig.**
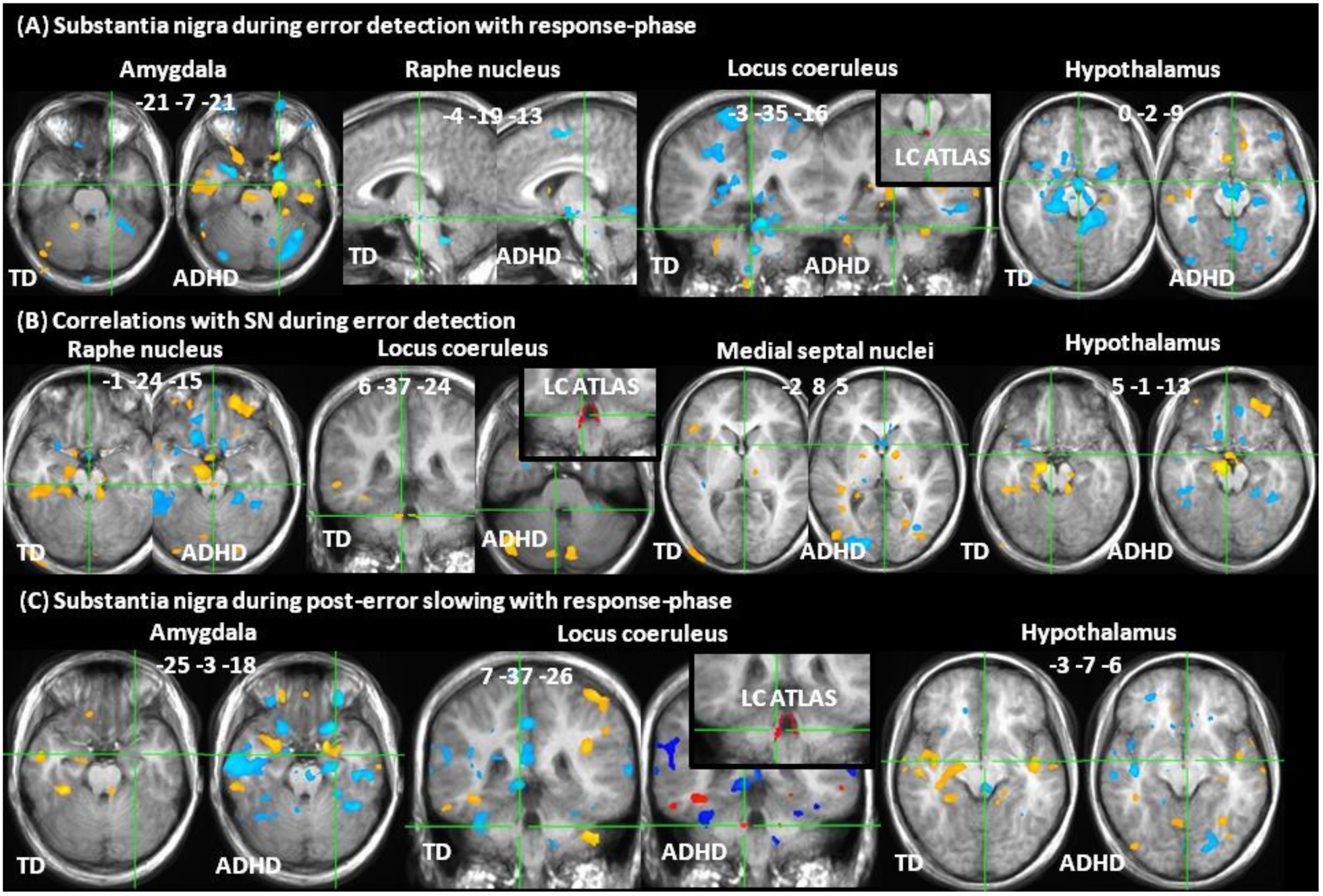
**Correlations with substantia nigra related to Table 3**. Correlation of substantia nigra during error detection with response-phase (A) and with error detection maps (B). Correlation of substantia nigra during post-error slowing with post-error slowing (C) and response-phase (D) maps. Correlation maps portray B1 estimates after whole brain correction (red/yellow = positive, blue = negative correlation). Locations in Talairach space, portrayed in radiological space (left = right); LC ATLAS = insets showing locus coeruleus atlas in red.

**S4 Fig.**
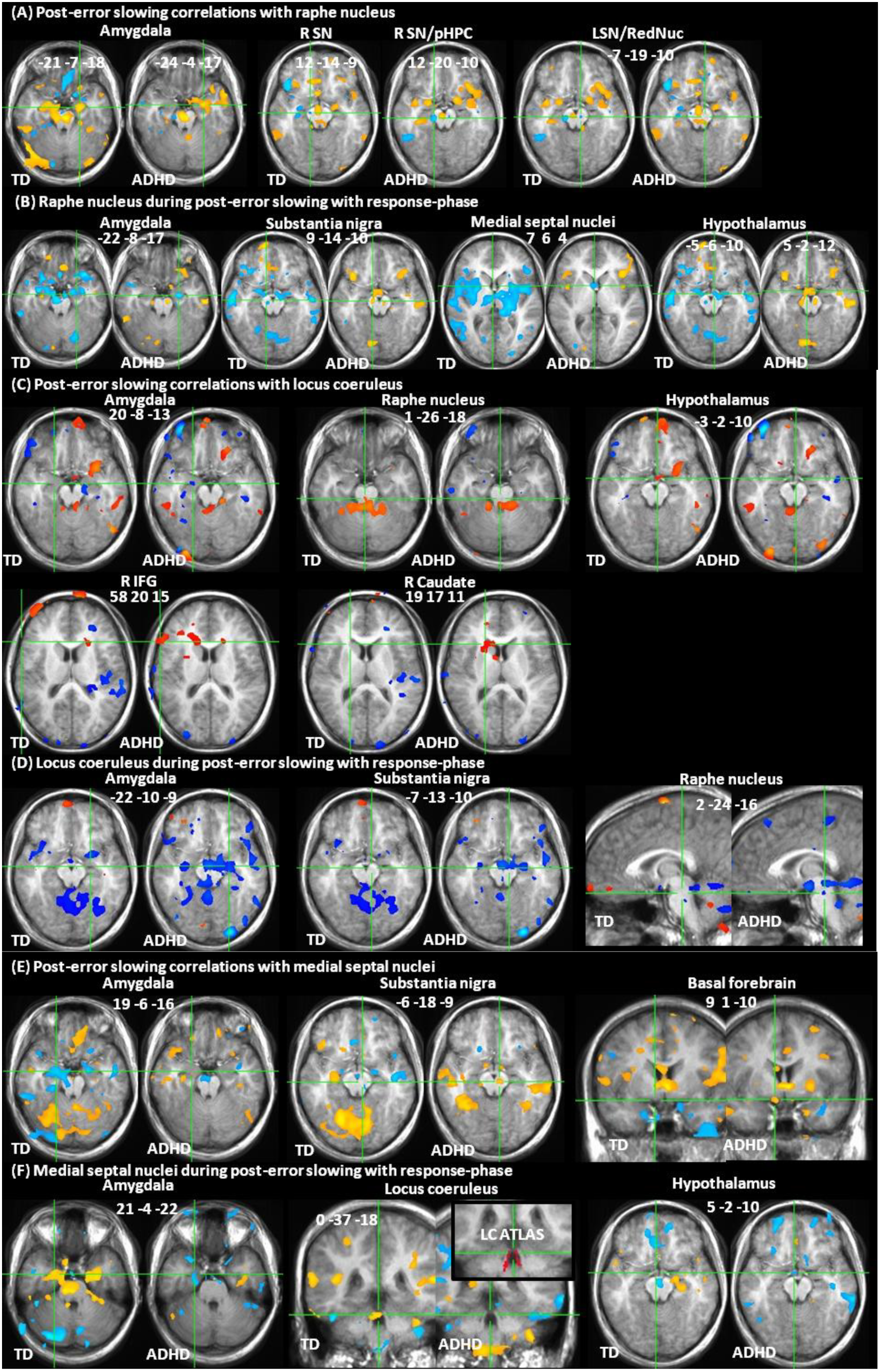
**Correlations with raphe nucleus from Table 4, with LC from Table 5, and with medial septal nuclei from Table 6.** (A) Correlation of raphe nucleus during post-error slowing with post-error slowing (A) and response-phase maps (B). Correlation of locus coeruleus activity during post-error with post-error slowing (C) and with response-phase maps (D). Correlation of medial septal nuclei during post-error slowing with post-error slowing (E) and response-phase maps (F). Correlation maps portray B1 estimates after whole brain correction (red/yellow = positive, blue =- negative correlation). Locations in Talairach space, portrayed in radiological space (left = right).

**S5 Fig.**
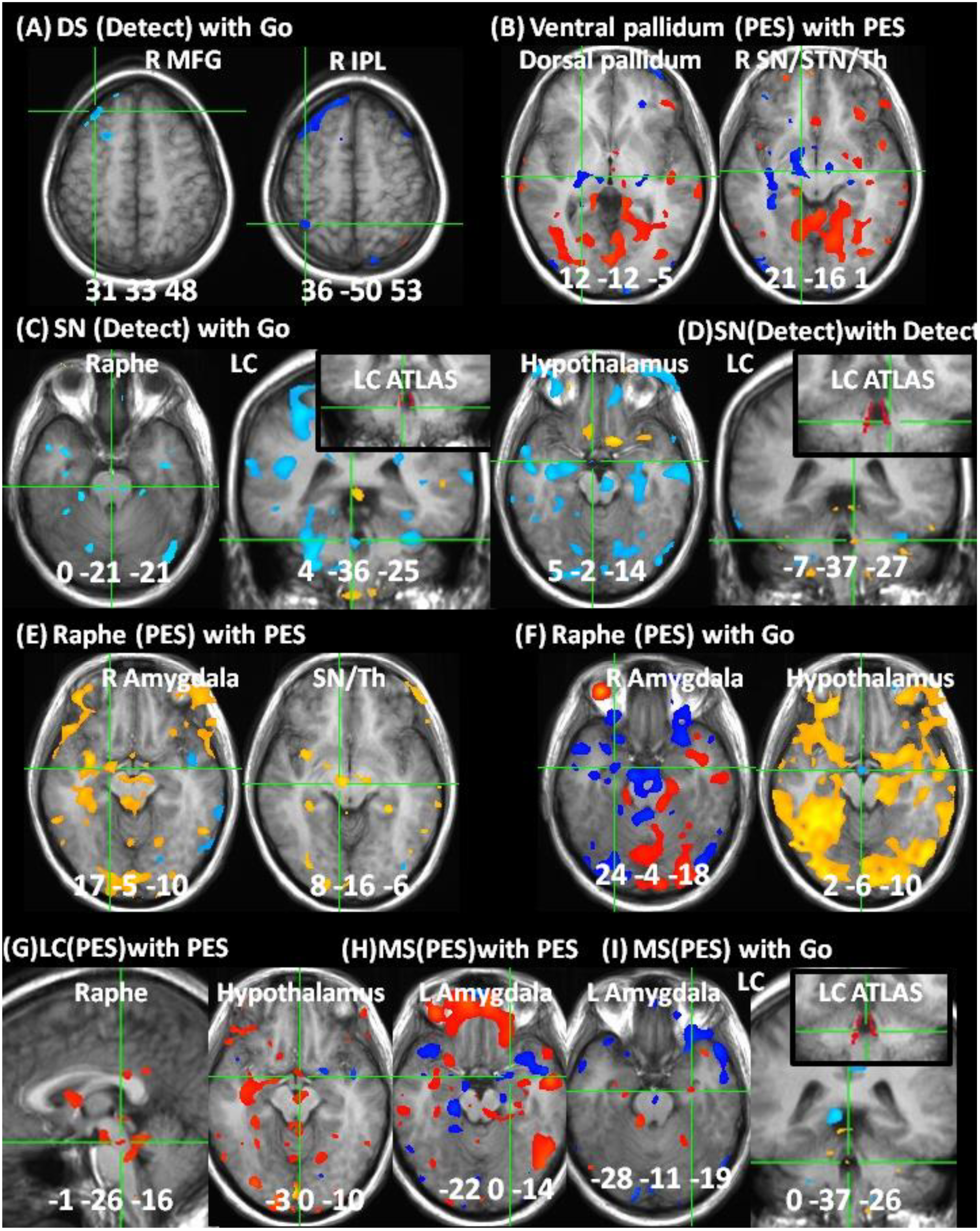
**Correlations in replication sample from Table 7.** (A) Correlation of dorsal striatum (DS) during error detection (Detect) with response-phase (Go) activities in right middle frontal gyrus (R MFG) and inferior parietal lobule (IPL). (B) Correlation of ventral pallidum with dorsal pallidum and substantia nigra (SN) during post-error slowing (PES). (C) Correlation of SN during Detect with Go activity in raphe nucleus, locus coeruleus (LC) and hypothalamus. (D) SN correlation with LC during Detect. (E) Raphe correlation with amygdala and SN during PES. (F) Correlation of raphe nucleus during PES with amygdala and hypothalamus during Go. (G) Correlation of LC with raphe nucleus and hypothalamus during PES. (H) Correlation of medial septal nuclei (MS) with amygdala during PES. (I) Correlation of MS during PES with LC during Go. Correlation maps (B1 estimates) were whole-brain corrected except where subthreshold (B ii,F i, H). Color bars depict Z-score range. Locations in Talairach space, portrayed in radiological space (left = right).

**S6 Fig.**
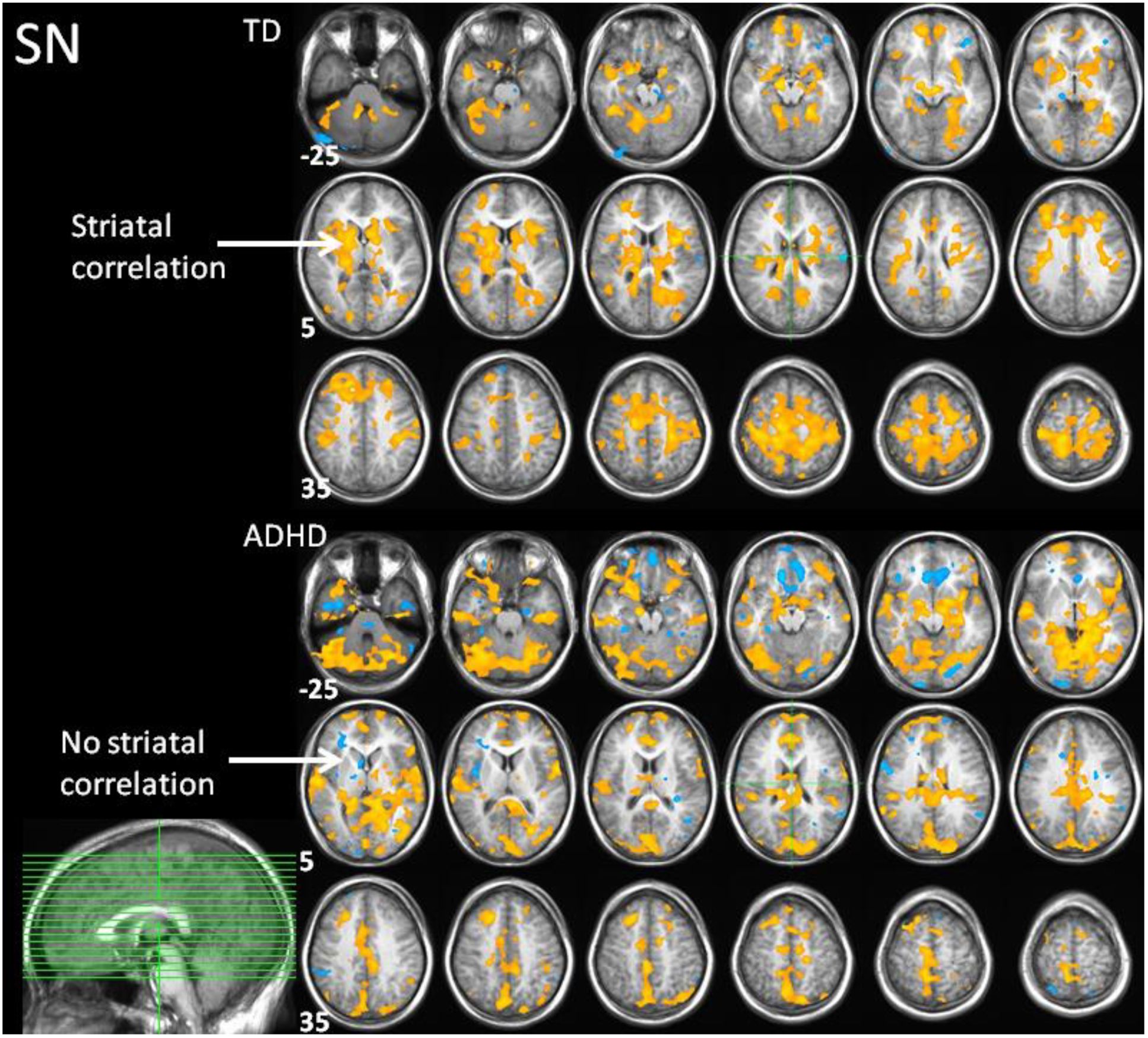
**Confirmatory analysis of SN seed location**. Whole brain corrected correlations (red/yellow = positive, blue = negative correlation) with SN seed during post-error slowing. In TD, SN correlated with bilateral limbic, striatal and neocortical regions, consistent with known ascending DA pathways. ADHD showed stronger correlations in posterior networks and negative correlations were present in rostral ACC. Striatal correlations were absent in ADHD, indicating a lack of nigrostriatal influence during post-error slowing. Numbers indicate slice locations in Talairach coordinates (ascending in 5mm increments), portrayed in midline sagittal plane at bottom left.

**S7 Fig.**
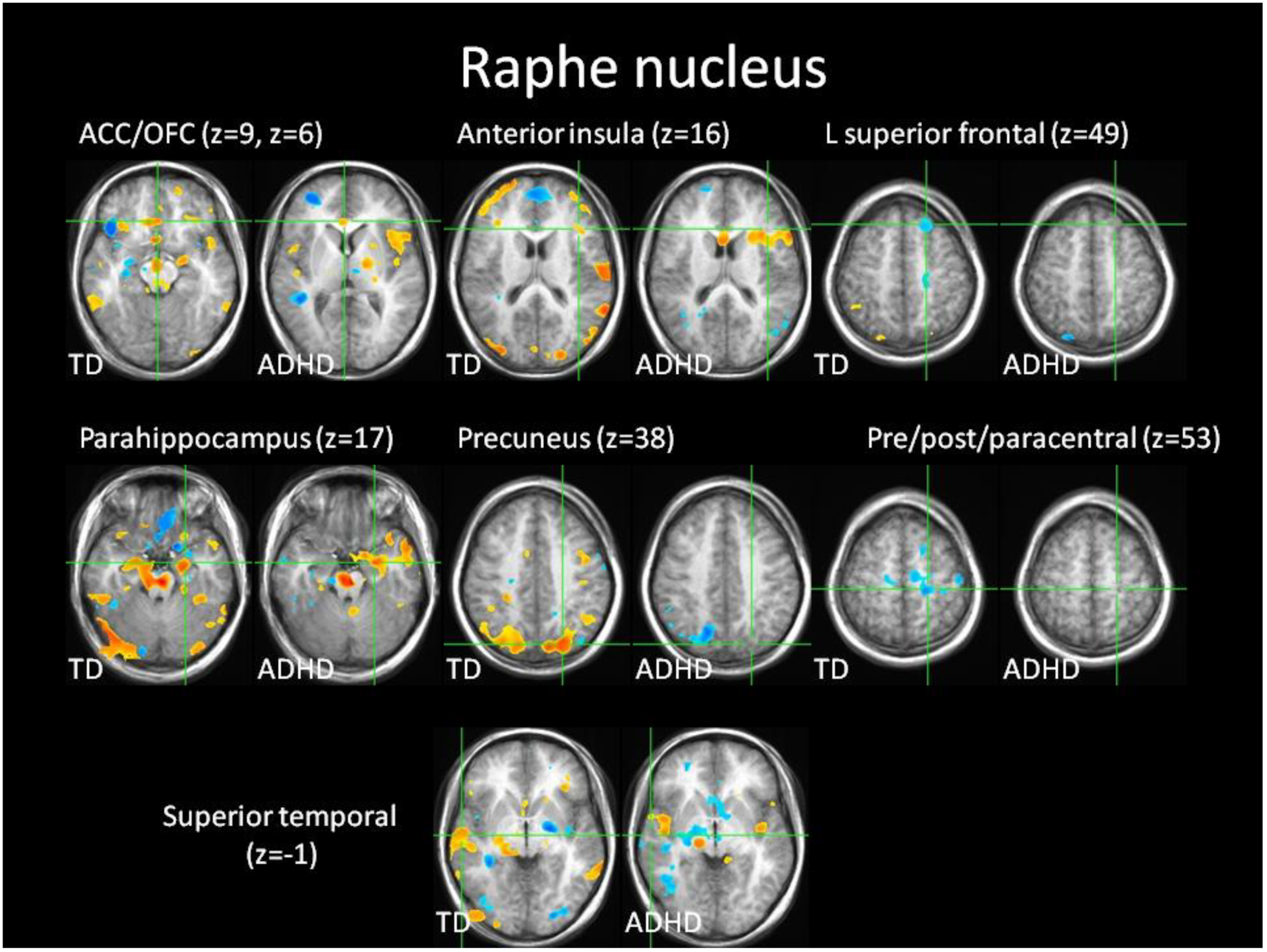
**Confirmatory analysis of raphe nucleus seed location.** Whole brain corrected correlations (red/yellow = positive, blue = negative correlation) with raphe nucleus seed during post-error slowing in the same regions and direction (positive/negative) as in (Beliveau et al., 2015). Correlations were weaker or absent in ADHD except in anterior insula, where correlations were stronger than in TD. z = slice location in Talairach coordinates.

